# Development, Characterization, and Structural Analysis of a Genetically Encoded Red Fluorescent Peroxynitrite Biosensor

**DOI:** 10.1101/2023.04.13.536636

**Authors:** Yu Pang, Mian Huang, Yichong Fan, Hsien-Wei Yeh, Ying Xiong, Ho Leung Ng, Hui-wang Ai

**Affiliations:** Center for Membrane and Cell Physiology, University of Virginia, Charlottesville, VA 22908, USA; Department of Chemistry, University of Virginia, Charlottesville, VA 22904, USA; Department of Biochemistry and Molecular Biophysics, Kansas State University, Manhattan, KS 66506, USA; Department of Molecular Physiology and Biological Physics, University of Virginia School of Medicine, Charlottesville, VA 22908, USA; The UVA Comprehensive Cancer Center, University of Virginia, Charlottesville, VA 22908, USA

## Abstract

Boronic acid-containing fluorescent molecules have been widely used to sense hydrogen peroxide and peroxynitrite, which are important reactive oxygen and nitrogen species in biological systems. However, it has been challenging to gain specificity. Our previous studies developed genetically encoded, green fluorescent peroxynitrite biosensors by genetically incorporating a boronic acid-containing noncanonical amino acid (ncAA), *p*-boronophenylalanine (*p*BoF), into the chromophore of circularly permuted green fluorescent proteins (cpGFPs). In this work, we introduced *p*BoF to amino acid residues spatially close to the chromophore of an enhanced circularly permuted red fluorescent protein (ecpApple). Our effort has resulted in two responsive ecpApple mutants: one bestows reactivity toward both peroxynitrite and hydrogen peroxide, while the other, namely pnRFP, is a selective red fluorescent peroxynitrite biosensor. We characterized pnRFP *in vitro* and in live mammalian cells. We further studied the structure and sensing mechanism of pnRFP using X-ray crystallography, ^11^B-NMR, and computational methods. The boron atom in pnRFP adopts an sp^2^-hybridization geometry in a hydrophobic pocket, and the reaction of pnRFP with peroxynitrite generates a product with a twisted chromophore, corroborating the observed “turn-off” fluorescence response. Thus, this study extends the color palette of genetically encoded peroxynitrite biosensors, provides insight into the response mechanism of the new biosensor, and demonstrates the versatility of using protein scaffolds to modulate chemoreactivity.

## INTRODUCTION

Peroxynitrite (ONOO^−^) is a highly reactive molecule formed by the reaction between nitric oxide (•NO) and superoxide anion (O_2_^•–^) via a diffusion-controlled process ^1,2^. The decomposition of peroxynitrite can further produce secondary radicals ^3,4^. In biological systems, peroxynitrite and its derived radicals act through direct or radical-mediated oxidation and nitration of biomolecules ^5-8^. In particular, tyrosine nitration, which is the most representative post-translational protein modification caused by peroxynitrite, has been considered a maker of nitrosative stress ^9^. Peroxynitrite is thus recognized as an important pathogenic mediator for various diseases, such as cardiovascular diseases, neurodegeneration, inflammation, and cancer ^10-12^. Additionally, low levels of peroxynitrite may play signaling roles ^13,14^. Furthermore, peroxynitrite has been identified as a potent cytotoxic effector of macrophages, and due to its cell permeability, macrophage-induced peroxynitrite can kill surrounding target cells or invading pathogens^15,16^

To better understand the pathophysiology of peroxynitrite and develop related therapies, it is crucial to track peroxynitrite in living systems ^17^. However, this is challenging due to peroxynitrite’s high reactivity, low steady-state concentration, and complex diffusion and reaction pathways in the cellular milieu. Fluorescent sensors have emerged as a promising approach for peroxynitrite detection, as they are sensitive, provide good signal-to-background ratios, and can be used with widely available fluorescence microscopy platforms ^18-21^. However, these sensors often exhibit cross-reactivity with other reactive oxygen and nitrogen species (ROS/RNS), such as hydroxy radical (•OH), hypochlorous acid (ClO^−^), and hydrogen peroxide (H_2_O_2_). Developing specific sensors for peroxynitrite remains a highly sought-after goal.

Fluorescent molecules containing boronic acid have been commonly used for detecting hydrogen peroxide and peroxynitrite ^22,23^. However, it is difficult to differentiate between the two using these molecules, because although arylboronic acid reacts with peroxynitrite more rapidly than hydrogen peroxide, hydrogen peroxide is more prevalent and abundant in biological systems. To address this issue, previous studies have created several boronic acid-based sensors that utilize N-B interactions to convert boronic acid into an sp^3^-hybridized boron species, which reduces reactivity with hydrogen peroxide and increases specificity for peroxynitrite ^19,24-26^. In particular, we previously created genetically encoded fluorescent peroxynitrite sensors, such as pnGFP and pnGFP-Ultra, by introducing a noncanonical amino acid (ncAA), *p*-boronophenylalanine (*p*BoF), into the chromophores of circularly permuted fluorescent proteins (cpFPs) ^24-26^. We utilized a genetic code expansion technology ^27^, which involves the expression of an engineered orthogonal tRNA and aminoacyl-tRNA synthetase pair, to incorporate ncAAs into proteins in a site-specific manner. cpFPs were employed in our studies due to their more accessible chromophores than fluorescent proteins (FPs) in the wild-type topology ^28^. The peroxynitrite-induced oxidation of boronic acid-derived chromophores generates tyrosine-derived phenolate chromophores (**Figure 1a**), resulting in drastic fluorescence turn-on responses. The genetic encodability of these sensors makes them compatible with directed evolution for response and specificity optimization ^26^, and it further allows for convenient dissemination and broad adaptation. In addition, signal sequences can be used to readily express the sensors in subcellular localizations ^28^. Mechanistic studies suggest that the boron atoms in these sensors are converted to be sp^3^-hybridized due to the N-B interaction with a nearby histidine residue through a polarized water bridge, leading to high specificity that can differentiate peroxynitrite from hydrogen peroxide ^25^.

**Figure 1.**
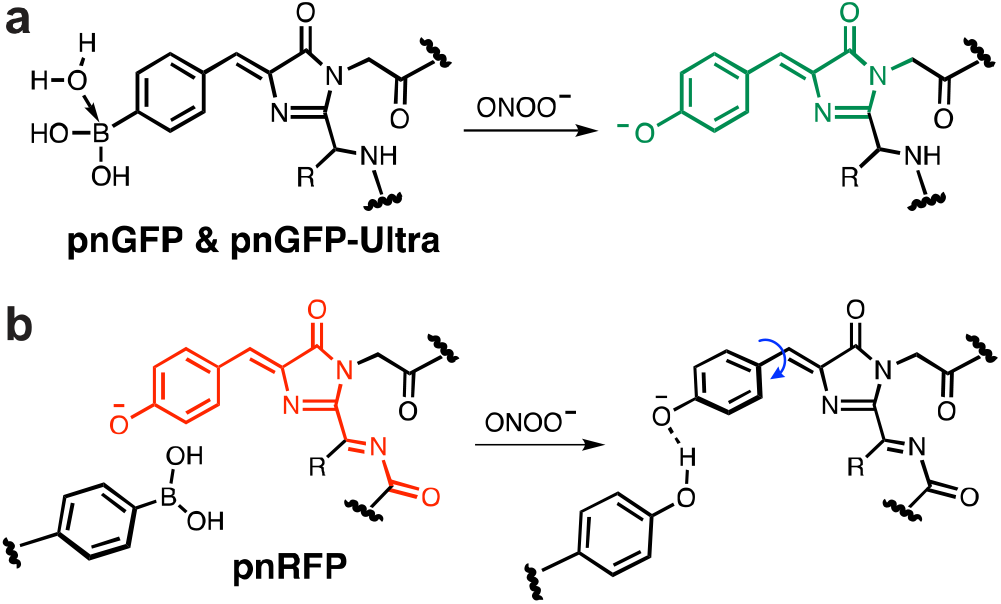
Mechanisms of protein-based, noncanonical amino acid (ncAA)-containing peroxynitrite biosensors. (**a**) **Previous work (pnGFP & pnGFP-Ultra):** *p*BoF was introduced to the chromophore of pnGFP or pnGFP-Ultra, which could further react with peroxynitrite to form a tyrosine-derived chromophore for enhanced fluorescence. (**b**) **This work (pnRFP):** *p*BoF is introduced to an amino acid residue in proximity to the chromophore, and the reaction of *p*BoF with peroxynitrite forms a tyrosine residue, which reduces fluorescence by bending the chromophore via hydrogen bonding.

Herein, we present the development, characterization, and structural analysis of a genetically encoded red fluorescent peroxynitrite biosensor (pnRFP), which specifically detects peroxynitrite *in vitro* and in mammalian cells. Instead of incorporating *p*BoF into the chromophore of a circularly permutated red fluorescent protein (cpRFP), we replaced residues near the chromophore with *p*BoF, allowing modulation of chromophore fluorescence through the boronic acid oxidation reaction (**Figure 1b**). In addition, we studied the structures of pnRFP and relevant mutants using biophysical and computational methods, gaining insight into the response mechanism of this new sensor.

## RESULTS

### Engineering of pnRFP

To expand the color palette of genetically encoded peroxynitrite biosensors, we sought to introduce *p*BoF into cpRFPs. We selected cpmApple, a cpRFP variant previously used in the development of other biosensors ^29,30^, as the starting scaffold. Next, we used error-prone polymerase chain reactions (EP-PCRs) to randomize cpmApple. Screening the libraries for increased brightness at 37 °C led to the identification of an enhanced cpmApple mutant (ecpApple), which is three mutations (S9G, E135K, and M206V) away from cpmApple (**Figure S1**). Next, taking inspiration from pnGFP and pnGFP-Ultra, we introduced *p*BoF to the chromophore-forming tyrosine residue of ecpApple via genetic code expansion. The codon of residue 176 of ecpApple was mutated to TAG (amber codon), and an amber suppression plasmid, pEvol-*p*BoF, which expresses orthogonal, *p*BoF-specific tRNA and aminoacyl-tRNA synthetase in *E. coli*, was used to incorporate *p*BoF in response to the amber codon ^31,32^. Unfortunately, our prepared protein (ecpApple-Y176B) did not show robust fluorescence responses to peroxynitrite. After mixing peroxynitrite with ecpApple-Y176B, we observed a very slow development of red fluorescence. Because the chromophore maturation of red fluorescent proteins (RFPs) has to undergo a more complex process than green fluorescent proteins (GFPs) ^33^, we reasoned that the introduction of *p*BoF to residue 176 of ecpApple disrupted the formation of a mature chromophore. We further performed random mutagenesis on ecpApple-Y176B but were unable to identify any improved mutants.

Because the fluorescence of FP chromophores is sensitive to the surrounding environment, we next investigated the possibility of replacing residues near the chromophore of ecpApple with *p*BoF. We selected residues 14, 28, 30, 48 and 66 (**Figure S2a**) and introduced *p*BoF to each of these residues. The variants were expressed and purified from *E. coli* and tested for fluorescence responses to 100 μM peroxynitrite (**Figure 2a**). To our delight, two of these variants, including ecpApple-S14B and ecpApple-K30B, showed turn-on and turn-off responses, respectively. The results suggest that the peroxynitrite-induced oxidation of the two residues, which are spatially close to the phenolate of the chromophore (**Figure S2b**), can modulate the fluorescence of the chromophore.

**Figure 2.**
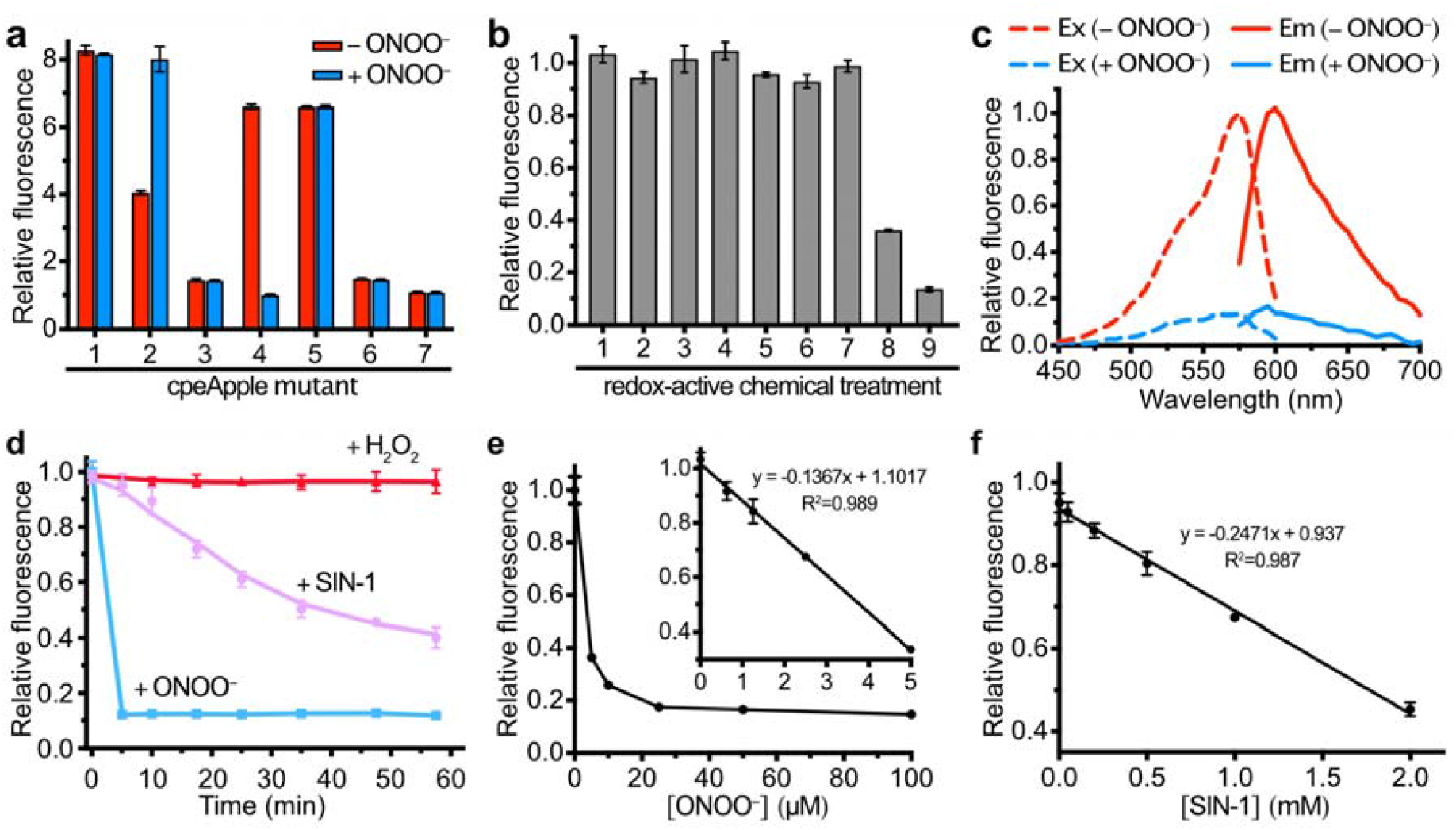
*In vitro* characterization of pnRFP and relevant mutants. (**a**) Relative fluorescence intensities of pnRFP and other relevant variants (1, ecpApple; 2, ecpApple-S14B; 3, ecpApple-I28B; 4, pnRFP (*a*.*k*.*a*. ecpApple-K30B); 5, ecpApple-Y48B; 6, ecpApple-L66B; and 7, pnRFP-B30Y (*a*.*k*.*a*. ecpApple-K30Y) before (red) and after (blue) treatment with 100 µM peroxynitrite. (**b**) Fluorescence responses of pnRFP after 20-min incubation with a series of redox-active chemicals: 1, PBS; 2, 5 mM GSH; 3, 5 mM L-cysteine; 4, 1 mM DTT; 5, 1 mM H_2_O_2_; 6, 300 μM O_2_^•−^; 7, 100 μM NaHS (H_2_S donor); 8, 5 μM ONOO^−^; 9, 100 μM ONOO^−^. (**c**) Excitation (dash line) and emission (solid line) spectra of pnRFP before (red) and after (blue) reaction with 100 μM ONOO^−^. (**d**) Time-dependent responses of pnRFP to 100 μM ONOO^−^ (blue), 10 mM SIN-1 (magenta), and 100 μM H_2_O_2_ (red). (**e**) Dose-dependent response of pnRFP (1 µM) to ONOO^−^ at indicated concentrations. (**f**) Dose-dependent response of pnRFP (1 µM) to SIN-1 at indicated concentrations. The incubation was 1 h at room temperature. Represented spectra are shown in panel c, and data in all other panels are presented as mean ± s.d. of triplicates.

We further tested ecpApple-S14B and ecpApple-K30B in response to hydrogen peroxide. The fluorescence of ecpApple-S14B was sensitive to micromolar hydrogen peroxide, and 100 μM hydrogen peroxide increased the fluorescence of ecpApple-S14B similarly to 100 μM peroxynitrite (**Figure S3**). In contrast, the fluorescence of ecpApple-K30B was unaffected by 1 mM hydrogen peroxide (**Figure 2b**). Due to the selectivity of ecpApple-K30B toward peroxynitrite, we named this mutant “pnRFP”. Furthermore, we constructed a pnRFP-B30Y mutant (equivalent to ecpApple-K30Y, **Figure S1**), which is the expected major oxidation product of pnRFP (**Figure 1a**). pnRFP-B30Y was unresponsive to peroxynitrite (**Figure 2a**), confirming that the quenching of pnRFP by peroxynitrite was indeed caused by the reaction between the *p*BoF30 residue and peroxynitrite.

### Further characterization of pnRFP *in vitro*

We examined the specificity of pnRFP against an expanded panel of redox-active molecules involved in cellular redox signaling. At physiologically relevant concentrations, only peroxynitrite caused notable fluorescence changes (**Figure 2b**), further confirming that pnRFP is a specific peroxynitrite sensor. pnRFP emitted strong red fluorescence with the excitation and emission peaks at 572 and 594 nm, respectively (**Figure 2c**). After reacting with peroxynitrite, its fluorescence decreased by about 5-fold. The reaction between pnRFP and peroxynitrite completed quickly, while a prolonged incubation of pnRFP with hydrogen peroxide did not cause notable fluorescence changes (**Figure 2d**). In addition, the incubation of pnRFP with SIN-1 (3-morpholinosyndnomine), a slow peroxynitrite-releasing molecule ^34^, resulted in a time-dependent fluorescence decrease (**Figure 2d**). Furthermore, we mixed pnRFP with various concentrations of peroxynitrite or SIN-1, and the induced fluorescence changes were concentration-dependent (**Figure 2e,f**). In particular, pnRFP was quite sensitive to peroxynitrite, and the response was saturated by ∼5 µM peroxynitrite. A linear response was observed in the low micromolar concentration range (< 5 µM) with a limit of detection (LOD) of ∼225 nM. The linear response range of pnRFP for SIN-1 was in the low millimolar range (< 2 mM). Collectively, the results suggest that pnRFP is a specific and sensitive red fluorescent peroxynitrite sensor.

### Use of pnRFP to image peroxynitrite in live mammalian cells

To examine whether pnRFP could image peroxynitrite in mammalian cells, we co-transfected human embryonic kidney (HEK) 293T cells with our constructed mammalian expression plasmid pMAH-pnRFP (**Figure S4a**), in addition to pMAH-POLY-eRF1(E55D) ^26^ which expresses an orthogonal tRNA and aminoacyl-tRNA synthetase pair and a translation release factor mutant for efficient genetic encoding of *p*BoF in mammalian cells (**Figure S4b**). Cells were treated with the peroxynitrite donor SIN-1, and the fluorescence of pnRFP-expressing cells decreased as peroxynitrite was generated, causing a 22% (∆F/F_0_) fluorescence decay within the monitoring period (**Figure 3a-c**). pnRFP-B30Y, whose fluorescence is insensitive to peroxynitrite, was expressed as a negative control. HEK 293T cells expressing pnRFP-B30Y showed negligible fluorescence changes under the SIN-1 treatment. Next, we expressed pnRFP in mouse RAW264.7 macrophage cells. A plasmid (pAcBAC3-POLY-pnRFP, **Figure S4c**) containing the genes for pnRFP, the engineered aminoacyl-tRNA synthetase, and 20 copies of the amber suppression tRNA was used to boost protein expression. After transfection and pnRFP expression, RAW264.7 cells were pre-treated with lipopolysaccharide (LPS) and interferon-γ (INF-γ) to induce the expression of nitric oxide synthase. Next, phorbol 12-myristate 13-acetate (PMA) was added to activate NADPH oxidase, and cells were imaged simultaneously. We observed a 20% (∆F/F_0_) fluorescence turn-off response in these activated pnRFP-expressing cells, while cells expressing the negative control, pnRFP-B30Y, showed no obvious fluorescence change (**Figure 3d-f**). Collectively, these results confirm that pnRFP is a reliable biosensor for monitoring chemically induced and physiologically relevant peroxynitrite generation in live mammalian cells.

**Figure 3.**
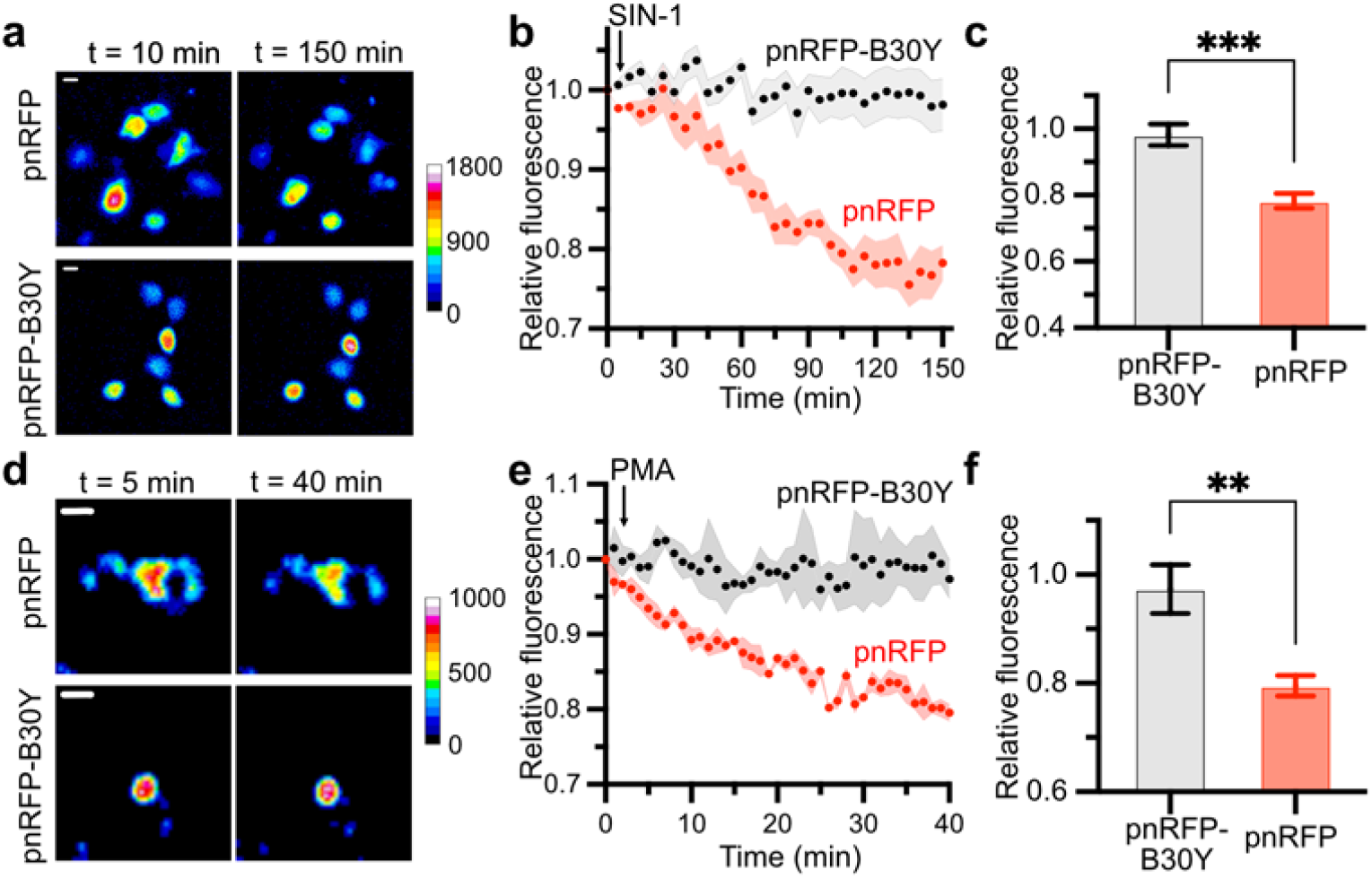
Imaging peroxynitrite in mammalian cells using pnRFP. (**a**) Representative pseudocolored fluorescence images of pnRFP- or pnRFP-B30Y-expressing HEK 293T cells in response to SIN-1 (1 mM). Images were taken at 10 and 150 min. Scale bar, 10 µm. (**b**) Time-lapse quantitation of fluorescence changes of HEK 293T cells expressing either pnRFP (red) or pnRFP-B30Y (black) in response to SIN-1. Intensities were normalized to the fluorescence signal of each cell at *t* = 0 min. Arrow indicates the time point for SIN-1 addition. (**c**) Comparison of total intensity changes for HEK 293T cells expressing pnRFP (red) or pnRFP-B30Y (black). Data represent mean ± s.e.m. of 9 cells from three technical repeats in each group (\*\*\**P* < 0.001, unpaired two-tailed *t*-test). (**d**) Representative pseudocolored fluorescence images of pnRFP- or pnRFP-B30Y-expressing RAW 264.7 cells in response to PMA (100 nM/mL). Images were taken at 5- and 40-min. Scale bar, 10 µm. (**e**) Time-lapse quantitation of fluorescence changes of RAW 264.7 cells expressing either pnRFP (red) or pnRFP-B30Y (black) in response to treatment of PMA followed 15 hr incubation in LPS/IFN γ. Intensities were normalized to the fluorescence signal of each cell at *t* = 0 min. Arrow indicates the time point for PMA addition. (**f**) Comparison of total intensity changes for RAW 264.7 cells expressing pnRFP (red) or pnRFP-B30Y (black). Data represent mean ± s.e.m. of 3 cells from three technical repeats (\*\**P* < 0.01, unpaired two-tailed *t*-test).

### Structural and mechanistic analysis

To gain more insights into the sensing mechanism of pnRFP, we obtained the monomeric crystal structure of pnRFP (**PDB 7LQO**) to 2.10-Å resolution through X-ray crystallography (**Table S1**). The structure displayed the typical architecture of a fluorescent protein, in which a chromophore was surrounded by an 11-β-strand barrel (**Figure 4a**). In the crystal structure, the chromophore (denoted as NRQ176) formed from M175, Y176, and G177 (**Figure S1**) perfectly fitted in the 2Fo-Fc electron density map, indicating a clear *cis* conformation (**Figure S5**). Its approximately planar conformation was stabilized by the joint effects of hydrogen bonds and hydrophobic interactions (**Figure 4b**). Two carbonyl oxygen atoms on the backbone and the imidazole ring of NRQ176 contacted with Q173, K179, R204, Q218, and S220 via direct or water-mediated hydrogen bonds. In the meantime, other surrounding residues, including *p*BoF30, hydrophobically interacted with NRQ176.

**Figure 4.**
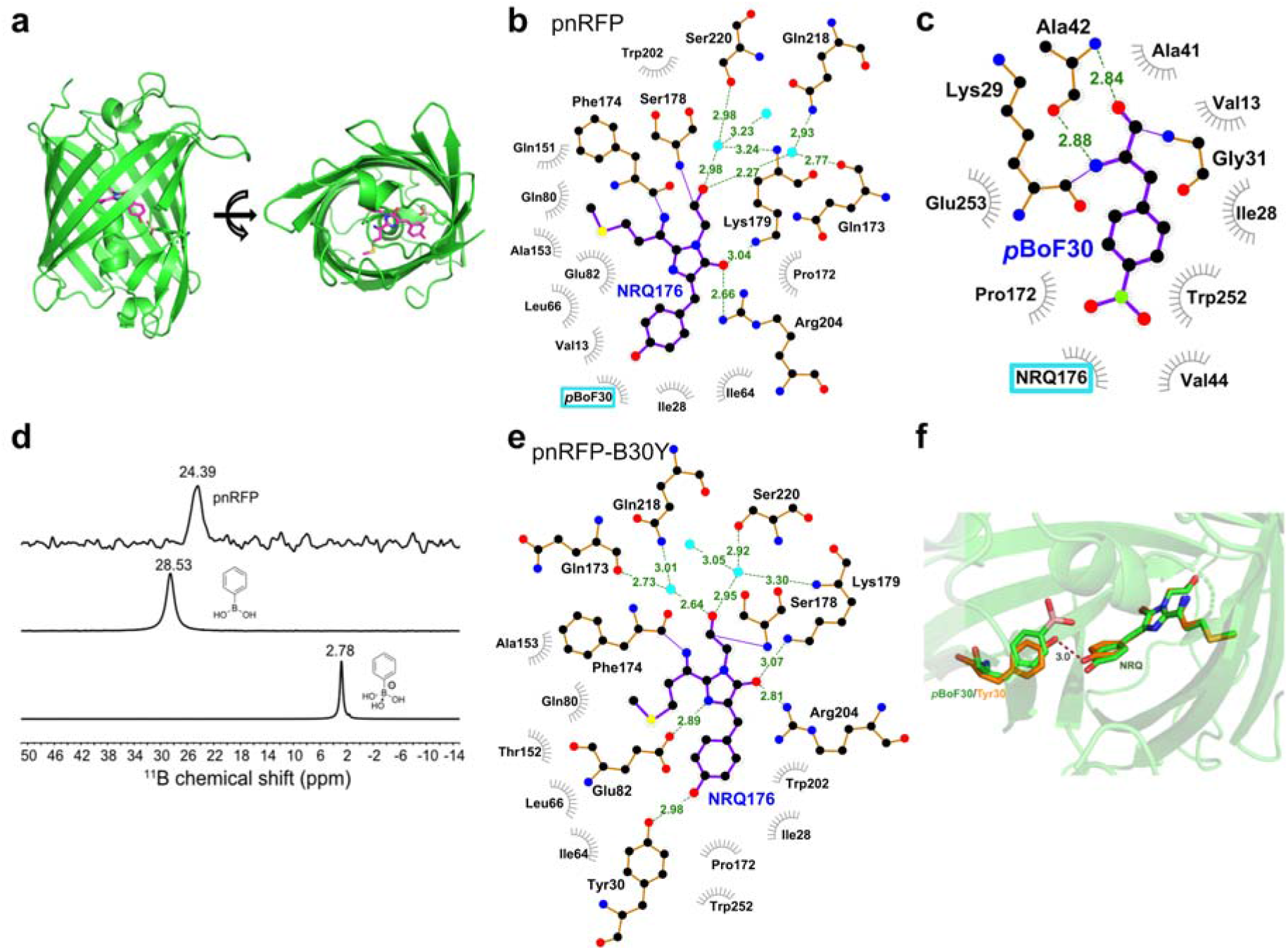
Structural analysis of pnRFP. (**a**) Side (left) and top (right) views of the X-ray crystal structure of pnRFP at 2.10 Å resolution (**PDB 7LQO**). (**b**) Interactions of the chromophore (NRQ176, purple) of pnRFP with surrounding residues through hydrogen bonds (green dash lines) and hydrophobic effects (red scattered lines). The conformation of the phenolic ring of the chromophore is stabilized by hydrophobic interactions with the surrounding residues, including *p*BoF30. (**c**) Interactions of *p*BoF30 in pnRFP with surrounding residues through hydrogen bonds (green dash lines) and hydrophobic effects (red scattered lines). (**d**) ^11^B-NMR spectra (from top to bottom: pnRFP, phenylboronic acid, and phenylboronic acid in 1 M NaOH), confirming an sp^2^-hybridized boron atom in pnRFP. (**e**) Interactions of the chromophore (NRQ176, purple) of the pnRFP-B30Y mutant (1.95 Å resolution, PDB ID 7LUG) with the surrounding residues, showing new hydrogen bonds including one between the phenolic ring of the chromophore and Tyr30. (**f**) Overlay of the structures of pnRFP (green) and the pnRFP-B30Y mutant (orange), highlighting out-of-plane distortion of the chromophore in pnRFP-B30Y.

The conformation of *p*BoF30 was described by the 2Fo-Fc electron density map at 1.0 σ (**Figure S6a**). To further confirm the conformation of the dihydroxyl boron portion, we replaced *p*BoF with Phe in the model by Coot ^35^ and recalculated electron density maps by Refmac5 ^36^. The positive Fo-Fc difference density appeared in the map (shown as an orange meshed bubble in **Figure S6b**), suggesting that we initially assigned the correct pose to *p*BoF30. The boron atom in pnRFP seemed to be sp^2^-hybridized, and the backbone of *p*BoF30 was stabilized through hydrogen bonding to A42, while the conformation of the benzene ring and the slightly twisted boronic acid head appeared to be an outcome of the hydrophobic interaction with the chromophore (NRQ176) and other residues (**Figure 4c**).

We further performed ^11^B-NMR spectroscopy to confirm that the sp^2^-hybridized boron in pnRFP was not an artifact caused by crystallization. The recorded spectrum of the purified pnRFP protein displayed a single peak at 24.39 ppm (**Figure 4d**). We also recorded the ^11^B-NMR spectra of phenylboronic acid and phenylboronic acid in 1 M NaOH. A single sharp peak of sp^2^-hybridized boron (free phenylboronic acid) was observed at 28.53 ppm, while sp^3^-hybridized boron (phenyl boronate anion) showed a peak at 2.78 ppm. Thus, the ^11^B-NMR spectra also support that the boron atom in pnRFP is sp^2^-hybridized, corroborating the X-ray crystallography result.

Next, we crystalized pnRFP-B30Y (*a*.*k*.*a*. ecpApple-K30Y, see **Figure S1**), which is the expected major oxidation product of pnRFP. The crystallization condition was identical to that used for pnRFP. The structure of pnRFP-B30Y, which was refined to a 1.95 Å resolution (**PDB 7LUG**), could superimpose on pnRFP with a 0.75 Å RMSD across Ca carbon atoms, showing a high degree of conformational similarity between the two proteins. In pnRFP-B30Y, two carbonyl oxygen atoms of the chromophore (NRQ176) connected to the surrounding residues through an almost identical hydrogen bonding network (**Figure 4e**). Meanwhile, two additional hydrogen bonds were formed between Y30 and the oxygen on the phenolic ring of the chromophore, and between Q82 and a nitrogen atom on the imidazole ring of the chromophore (**Figure 4e**). Three hydrogen bonds on the imidazole ring could limit the dynamics of the chromophore in space, while the phenolic rings of Y30 and the chromophore shifted closer to each other due to the new hydrogen bond between them. The combined forces disfavored the maintenance of the co-planar conformation of the chromophore. Subsequently, the bending of the chromophore (**Figure 4f**) reduced the conjugation effect through the π system, thereby leading to the diminished fluorescence of pnRFP-B30Y compared to pnRFP. Together, the structural results explain how peroxynitrite turns off the fluorescence of pnRFP.

Finally, we built a homology model for ecpApple-S14B, which was responsive to both peroxynitrite and hydrogen peroxide (**Figure S3**). We used the pnRFP structure (**PDB 7LQO**) as the scaffold and substituted *p*BoF30 with K with an extended sidechain conformation towards the phenolic ring of the chromophore. We also substituted S14 with *p*BoF in Coot ^35^ with the side chain buried in the β-barrel. Next, the model was then subjected to a 131-ns length molecular dynamics (MD) simulation with the AMBER14 force field in YASARA (**Figure S7**) ^37,38^. The average model indicated the dominant conformations of residues during the simulation, so we used it for further analysis. In this model (**Figure 5**), ecpApple-S14B well maintained a typical β-barrel structure resembling other fluorescent proteins. Also, *p*BoF14 π-stacks with the phenolic ring of the chromophore, and K30 interacts with the backbone of *p*BoF14 through a hydrogen bond. E82 further interacts with the hydroxyl group of *p*BoF14 through hydrogen bonding. Compared with *p*BoF30 in pnRFP (**Figure 4c**), the boronic acid-head group of *p*BoF14 in ecpApple-S14B stays in a more hydrophilic pocket (**Figure 5** and **Figure S8**), which favors the polarization of the group. Moreover, the negative charge carried by E82 drives electrons to move from *p*BoF14 to E82 through the hydrogen bond. We further used the AM1 semi-empirical method in YASARA to assign charges to the boron atoms, and the charges of the boron atoms in pnRFP and ecpApple-S14B were +0.3562 and +0.4998, respectively, supporting that that boron in pnRFP is less positively charged. Since the oxidation of phenylboronic acid started with the nucleophilic attack by anionic species (*e*.*g*., ONOO^−^ or HOO^−^) ^25^, the result is aligned with the observed, much-reduced reactivity of pnRFP, compared to ecpApple-S14B, with HOO^−^. Meanwhile, because ONOO^−^ is more electronegative than HOO^−^, ONOO^−^ can still retain good reactivity with pnRFP. Overall, the modeling results further confirm that protein scaffolds can offer unique local environments to modulate the reactivity of boronic acid.

**Figure 5.**
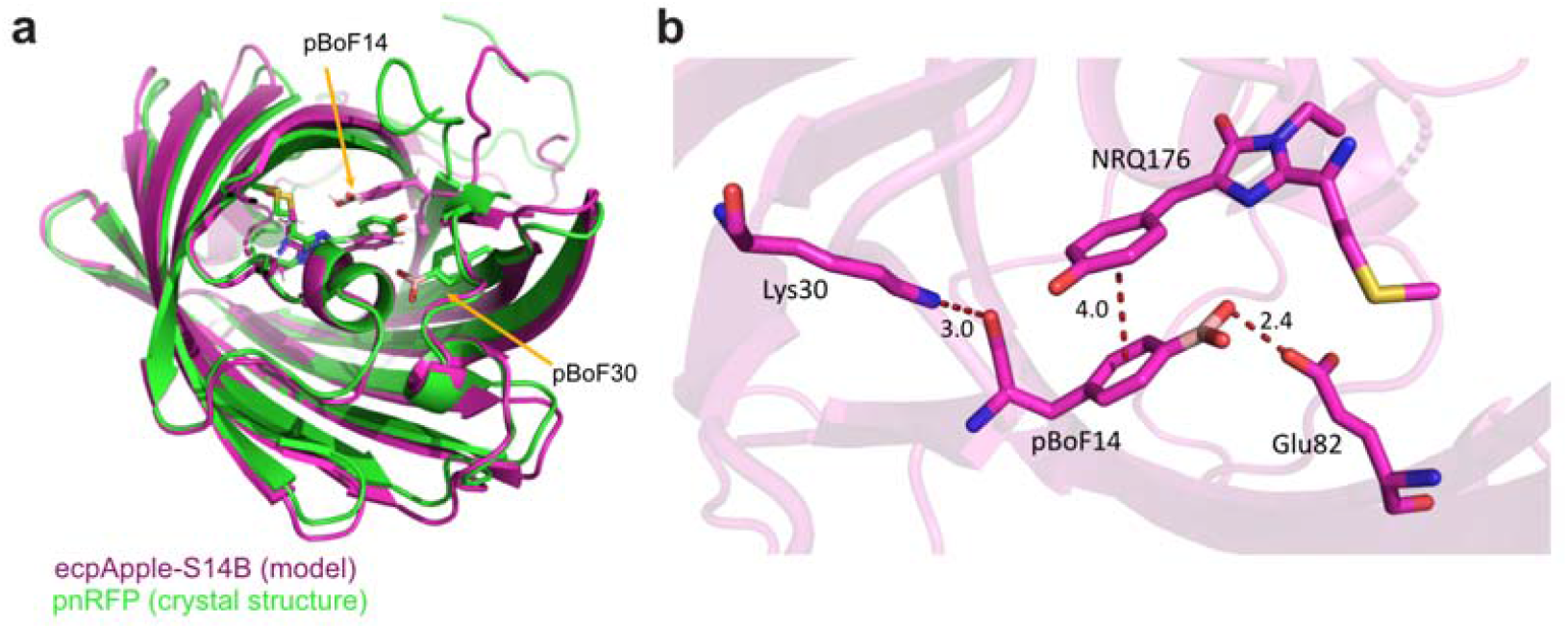
Computational modeling of ecpApple-S14B. (**a**) Overlay of the pnRFP crystal structure (green) and the model structure of ecpApple-S14B (magenta), highlighting the locations of *p*BoF30 and *p*BoF14 in relation to the chromophores. (**b**) Model structure of ecpApple-S14B, highlighting interactions (hydrogen bonds and π-π stacking) with *p*BoF14.

## DISCUSSION

Starting from a cpRFP mutant, we developed a genetically encoded red fluorescent peroxynitrite biosensor (pnRFP) by site-specifically introducing a boronic acid-containing ncAA into a residue close to the chromophore. This effort expands not only the color of genetically encoded peroxynitrite biosensors from green to red but also how ncAA-base FP biosensors could be designed. In addition to directly using ncAAs to modify the chromophores of FPs ^28,39-41^, modifying chromophore-surrounding residues with ncAAs has now proven to be another viable strategy for developing ncAA-base FP biosensors.

pnRFP were discovered from site-directed mutagenesis of a cpRFP mutant derived from the Ca^2+^ sensor, R-GECO1 ^29^. We introduced *p*BoF to several residues around the chromophore and gained responsive mutants at two locations (residues 14 and 30). This observation is not surprising, since the two residues have strong effects on the fluorescence of the chromophore. The crystal structure of R-GECO1 at the Ca^2+^-bound state was solved previously (**PDB 4I2Y**) ^42^. The K78 residue of R-GECO1 (equivalent to residue 30 in ecpApple or pnRFP) forms an ionic interaction with the phenolated oxygen of the chromophore, while S62 (equivalent to residue 14 in ecpApple or pnRFP) further stabilizes this interaction via a hydrogen bond (**Figure S2b**).

In pnRFP-B30Y, a hydrogen bond is formed between Y30 and the phenolated oxygen of the chromophore. A similar hydrogen bond exists between K78 and the chromophore of R-GECO1 at the Ca^2+^-bound state. However, the location of Y30 of pnRFP-B30Y has been shifted from that of K78 in R-GECO1 (**Figure S9**). The outcome is that K78 helps to maintain a largely co-planar conformation of the chromophore of R-GECO1, while Y30 contributes to bending the chromophore of pnRFP-B30Y. Therefore, R-GECO1 at the Ca^2+^-bound state is quite fluorescent, while pnRFP-B30Y shows reduced fluorescence.

Normally, arylboronic acid can be oxidized by both hydrogen peroxide and peroxynitrite to generate the phenol product. Previous studies used N-B interactions to generate sp^3^-hybridized boron species for enhanced specificity toward peroxynitrite ^19,24-26^. This study derived pnRFP, which still has an sp^2^-hybridized boron atom, as confirmed by the X-ray crystal structure and ^11^B-NMR spectra (**Figure 4**). Although further research is still needed to pinpoint the mechanism, our results here indicate that hydrophobicity and hydrogen bonding patterns in the local environment and the charge distribution of the boron atom may be factors contributing to the high specificity of pnRFP. Collectively, pnRFP and ecpApple-S14B, as well as pnGFP and pnGFP-Ultra, offer a series of examples that demonstrate the versatility of using protein scaffolds to tune chemoselectivity.

In summary, we have developed a genetically encoded red fluorescent peroxynitrite biosensor and characterized it *in vitro* and for imaging chemically induced and physiologically relevant peroxynitrite generation in live mammalian cells. We further used structural, biophysical, and computational methods to study the response mechanism of this new sensor. Our results may provide inspiration to further biosensor development and the turning of chemoreactivity.

## METHODS

### Sources of Key Reagents

The amino acid *p*-borono-DL-phenylalanine (*p*BoF) was purchased from Synthonix (Wake Forest, NC). Synthetic DNA oligos were purchased from Integrated DNA Technologies (Coralville, IA). Restriction endonucleases or other molecular biology reagents were purchased from Thermo Fisher Scientific (Waltham, MA) or New England Biolabs (Ipswitch, MA). Plasmid pCMV-R-GECO1 (Addgene Plasmid # 32444) was a gift from Robert E. Campbell.

### Mutagenesis and Biosensor Engineering

The generation of ecpApple from R-GECO1 was described elsewhere ^43^. Next, overlap extension polymerase chain reactions (PCRs) were used to introduce the TAG amber codon to specific residue positions of ecpApple. To introduce the TAG codon to residue 14, oligos ecpApple-S14TAG-F and cpRFP_R (**Table S2**) were first used to amplify a fragment from ecpApple; next, oligos cpRFP_F and cpRFP_R were used to further amplify and extend the fragment recovered from the previous reaction. The PCR product was then digested with Hind III and Xho I and ligated with a predigested, compatible pBAD vector. To introduce the TAG codon to residue 30, oligos cpRFP_F and ecpApple-K30TAG-R, and ecpApple-K30TAG-F and cpRFP_R were used in two separate PCRs to amplify two separate fragments from ecpApple. The two fragments were used as templates in an overlap extension PCR with oligos cpRFP_F and cpRFP_R. The full-length product was generated, digested with Hind III and Xho I, and inserted into a pBAD vector. A similar procedure was used to derive other ecpApple mutants, and the sequences of the corresponding oligos are presented in **Table S2**. Ligation products were used to transform *Escherichia coli* (*E. coli*) DH10B competent cell, which was next plated on LB agar plates supplemented with ampicillin (100 μg/mL). Plasmids were minipreped from liquid cell culture inoculated with single colonies, and the sequences of the variants were confirmed with Sanger Sequencing (Eurofins Genomics, Louisville, KY).

### Protein Purification and Characterization

The pBAD plasmid harboring the gene of each ecpApple mutant was used along with the pEvol-*p*BoF plasmid ^31,32^ to con-transform *E. coli* DH10B cells, which were next grown on LB agar plates supplemented with ampicillin (100 μg/mL), chloramphenicol (50 μg/mL), and L-arabinose (0.02% w/v) at 37 °C overnight. A single colony was used to inoculate 2×YT liquid culture with 100 μg/mL ampicillin and 50 μg/mL chloramphenicol. After growth overnight at 37 °C and 250 rpm, the saturated cell culture was diluted 100-fold with 2×YT with 100 μg/mL ampicillin and 50 μg/mL chloramphenicol. When the optical density at 600 nm (OD_600_) reached 0.6, 0.2% (w/v) L-arabinose and 2 mM racemic *p*BoF were added. Cells were grown at 30°C, 250 rpm for another 48 h, before being harvested by centrifugation and lysed by sonication. His-tagged proteins were purified using Pierce Ni-NTA agarose beads according to the manufacturer’s instructions. The buffer was switched to 1× phosphate-buffered saline (PBS, pH 7.4) via dialysis, and protein concentrations were determined with Bradford assays. To prepare in vitro assays, peroxynitrite was freshly prepared and diluted with 0.3 M ice-cold NaOH, while SIN-1 was freshly dissolved in DMSO. The preparation of other redox-active species was performed as described ^24^, and 1× PBS was used to dilute proteins and reagents. To record fluorescence excitation spectra, the emission wavelength was set at 620 nm, and the excitation was scanned from 450 to 600 nm. To record the emission spectra, the excitation was set at 540 nm, and the emission was scanned from 560 to 700 nm. Other intensity measurements typically used 560 nm excitation and 610 nm emission. To examine responses to various redox-active chemicals, protein stock solutions were diluted with 1× PBS to gain a final concentration of 1 μM. Individual redox-active molecules (5 μL) were added to proteins (95 μL) in individual wells of a 96-well plate sitting on ice. The mixtures were moved to room temperature and incubated for 20 min before fluorescence recording. To examine time-dependent responses, 95 μL of the protein in 1× PBS was mixed with 5 μL of each reagent, and the kinetics was monitored at room temperature for 1 h. To test concentration-dependent responses, the pnRFP protein was mixed with peroxynitrite or SIN-1 at the indicated concentrations in individual wells of a black 96-well plate. The mixtures were incubated at room temperature for 1 h before end-point measurements were performed. All fluorescence signals were recorded with a monochromator-based BioTek Synergy Mx Microplate Reader.

### Mammalian Expression and Live-Cell Imaging

To construct mammalian expression plasmids, oligos pnRFP_F and pnRFP_R were used to amplify the gene fragment of pnRFP or pnRFP-B30Y from corresponding pBAD plasmids. The resultant PCR product was digested with Hind III and Apa I and inserted into a pre-digested pMAH-POLY plasmid. The resultant pMAH-pnRFP or pMAH-pnRFP-B30Y plasmid was used along with pMAH-POLY-eRF1(E55D) to co-transfect human embryonic kidney (HEK) 293T cells by following a described procedure ^26^. The *p*BoF amino acid was added to the cell culture medium to a final concentration of 2 mM at 18 h after transfection. Next, the culture was kept in a 37 °C, 5% CO_2_ incubator for another 48 h, before being transferred into a fresh medium with no *p*BoF. Cells were imaged 16 h later in Dulbecco’s phosphate-buffered saline (DPBS) supplemented with 1 mM Ca^2+^ and Mg^2+^ on a Leica DMi8 inverted fluorescence microscope. To express pnRFP in mouse RAW 264.7 macrophage cells, oligos pBac3-F and pBac3-R were used to amplify the gene from pMAH-pnRFP, and the PCR product was inserted into a compatible pAcBac3 vector predigested with Xho I and EcoR I, resulting in pAcBAC3-POLY-pnRFP. The control plasmid pAcBAC3-POLY-pnRFP-B30Y was created similarly. RAW 264.7 cells were cultured in Dulbecco’s Modified Eagle Medium (DMEM) with GlutaMAX (Gibco) supplemented with 10% fetal bovine serum (FBS). 3 μg pAcBAC3 plasmid was used to transfect RAW 264.7 cells by mixing the DNA with 6 μg X-tremeGENE HP DNA Transfection Reagent (Roche) according to the manufacturer’s instruction. At 20 h after transfection, 1 mM *p*BoF was added dropwise. After another 24 h, the *p*BoF-containing medium was replaced with fresh DMEM with GlutaMAX and 10% FBS but no *p*BoF. After pBoF depletion, RAW 264.7 cells were incubated with 1 μg/mL LPS and 100 ng/mL IFN-γ for 15 h. Next, the culture media was replaced with a HEPES-buffered Hank’s Balanced Salt Solution (HBSS), and cells were treated with 100 nM/mL PMA and imaged on a Leica DMi8 inverted fluorescence microscope. A Leica EL6000 light source, a TRITC filter cube (545/25 nm bandpass excitation and 605/70 nm bandpass emission), and a Photometrics Prime 95B sCMOS camera were used for the imaging experiments.

### Protein Crystallization and Structure Determination

22 mg/mL of the purified pnRFP in 20 mM Tris, pH 8.0 was crystallized by sitting-drop method with vapor diffusion against the crystallization reagent (0.1 M Na_2_HPO_4_: citric acid pH 4.2, 40% (v/v) PEG 400). A liquid drop at pH 6.0 was prepared by mixing 0.6 μL of protein solution with 0.6 μL of the crystallization reagent. It was equilibrated against 400 μL of the crystallization reagent to accomplish crystallization. To obtain high-quality crystals, microseeding was applied under the same crystallization condition. The mutant pnRFP-B30Y was crystallized at 8 mg/mL under the same process mentioned above without microseeding operation. X-ray diffraction data for the pnRFP crystal was collected by the SIBYLS beamline of the Advanced Light Source at Lawrence Berkeley National Lab, and the pnRFP-K30Y crystal was collected by the beamline 23-ID-D of the Argonne Photon Source, Chicago. Both data were processed by X-ray Detector Software (XDS) ^44^ and reduced by POINTLESS ^45^, AIMLESS ^46^ and TRUNCATE ^47^ in the CCP4 suite. Molecular replacement was applied to the reduced data with Phaser ^48^ for building structure models. The models were then refined by Coot 0.8.9-pre EL ^35^ and Refmac5 ^36^. Crystal structures of pnRFP and pnRFP-B30Y were performed via the PyMOL Molecular Graphics System, Version 1.8 Schrödinger, LLC. The protein-ligand 2D interaction diagrams were built via LigPlot+ version 1.4.5. ^49^ The RMSD between the structures was calculated by YASARA ^37^.

### ^11^B-NMR Characterization

The pnRFP protein was purified as described above, concentrated using Amicon Ultra Centrifugal Filter Units (3000 Da cutoff), and exchanged into 20 mM phosphate (pH 7.4, D_2_O (v): H_2_O (v) = 1:1) to final concentrations of 15.6 mg/mL. ^11^B NMR spectra were acquired on a Varian VNMRS 600 spectrometer operating at 192.439 MHz using a 5 mm AutoXDB probe. Samples were placed in quartz NMR tubes. Protein spectra were collected with single pulse excitation (45 degree pulses were used, 90 degree pulse width = 13 µsec). The sweep width was 200 ppm and the total acquisition time was 24 h (scan = 1.4 million, delay before pulse = 10 msec, single FID acquisition time = 52 msec). To further remove background signal, back linear prediction within the MestReNova software was used (Method = Toeplitz, COEF = 32, Base points = 256, from 0 to 46). Data were then processed in the normal manner with 100 Hz line broadening applied. Chemical shifts were referenced to external 15% boron trifluoride etherate in CDCl_3_ (BF_3_·Et_2_O, δ = 0 ppm). For standard comparisons, we recorded ^11^B NMR spectra for both of phenylboronic acid (20 mM) in the 20 mM phosphate buffer (pH 7.4, D_2_O (v): H_2_O (v) = 1:1), and phenylboronic acid (20 mM) in 1N NaOH aqueous solution (D_2_O (v): H_2_O (v) = 1:1).

### Computational Modeling

The crystal structure of pnRFP was used as a template. At position 30, BoF was substituted by Lys with an extended side-chain conformation towards the phenyl ring of NRQ. At position 14, Ser was substituted by BoF. BoF was adjusted in COOT to adopt a conformation with the side chain buried in the barrel. The homology model was further conformationally adjusted by MD simulations The model was then subjected to a 131-ns length molecular dynamics (MD) simulation with the AMBER14 force field in YASARA version 19.12.14.L.64. The system was performed with explicit solvent in a cube box with a 60.83-Å side length, containing 6,437 water molecules. The default parameter settings were used in the MD run with the pressure at 1 bar, the temperature of 298 K, pH 7.4, 1-fs time steps, and snapshots taken at every 100-ps interval. The trajectory was analyzed, and the simulated model was considered in the equilibrium state. The partial atomic charges assigned to boron and other atoms were calculated by the AM1 semi-empirical methods ^50^.

## ASSOCIATED CONTENT

### Supporting Information

The Supporting Information is available free of charge on the ACS Publications website. Supplementary figures and tables (PDF)

## AUTHOR INFORMATION

### Author Contributions

H.A. and H.L.N. conceived and supervised the project. Y.P. Y.F. and Y.X. engineered the biosensor, prepared proteins, and characterized the biosensor and related variants. M.H. solved the crystal structures and performed computational modeling. H.W.Y. and J.F.E. carried out the ^11^B-NMR experiment. H.A., Y.P., M.H., and H.L.N wrote the manuscript.

### Notes

The authors declare no competing interest.

## ACKNOWLEDGMENT

We thank Dr. Jeff F. Ellena at the UVA Biomolecular Magnetic Resonance Facility for technical assistance in acquiring the ^11^B-NMR data. Research reported in this publication was supported by funding to H.A. (University of Virginia Start-up Package and NIH grants R01 GM129291, R01 DK122253, and RF1 AG077773), and H.L.N. (Intel Corp and Nvidia Corp.).

## Notes

### Competing Interest Statement

The authors have declared no competing interest.

## REFERENCES

1. Radi, R., eroxynitrite, a stealthy biological oxidant. J. Biol. Chem. 2013, 288 (37), 26464–26472.

2. Beckman, J. S.; Koppenol, W. H., Nitric oxide, superoxide, and peroxynitrite: The good, the bad, and the ugly. Am. J. Physiol. Cell Physiol. 1996, 271 (5), C1424–C1437.

3. Szabó, C.; Ischiropoulos, H.; Radi, R., Peroxynitrite: biochemistry, pathophysiology and development of therapeutics. Nat. Rev. Drug Discov. 2007, 6 (8), 662–680.

4. Beckman, J. S.; Beckman, T. W.; Chen, J.; Marshall, P. A.; Freeman, B. A., Apparent hydroxyl radical production by peroxynitrite: implications for endothelial injury from nitric oxide and superoxide. Proc. Natl. Acad. Sci. U. S. A. 1990, 87 (4), 1620–1624.

5. Radi, R., Protein tyrosine nitration: biochemical mechanisms and structural basis of functional effects. Acc. Chem. Res. 2013, 46 (2), 550–559.

6. Alvarez, B.; Ferrer-Sueta, G.; Freeman, B. A.; Radi, R., Kinetics of peroxynitrite reaction with amino acids and human serum albumin. J. Biol. Chem. 1999, 274 (2), 842–848.

7. Rubbo, H.; Radi, R.; Trujillo, M.; Telleri, R.; Kalyanaraman, B.; Barnes, S.; Kirk, M.; Freeman, B. A., Nitric oxide regulation of superoxide and peroxynitrite-dependent lipid peroxidation. Formation of novel nitrogen-containing oxidized lipid derivatives. J. Biol. Chem. 1994, 269 (42), 26066–26075.

8. Kennett, E. C.; Davies, M. J., Glycosaminoglycans are fragmented by hydroxyl, carbonate, and nitrogen dioxide radicals in a site-selective manner: implications for peroxynitrite-mediated damage at sites of inflammation. Free Radic. Biol. Med. 2009, 47 (4), 389–400.

9. Monteiro, H. P.; Arai, R. J.; Travassos, L. R., Protein tyrosine phosphorylation and protein tyrosine nitration in redox signaling. Antioxid. Redox Signal. 2008, 10 (5), 843–890.

10. Félétou, M.; Vanhoutte, P. M., Endothelial dysfunction: a multifaceted disorder (the Wiggers Award Lecture). Am. J. Physiol. Heart Circ. 2006, 291 (3), H985–H1002.

11. Baillet, A.; Chanteperdrix, V.; Trocmé, C.; Casez, P.; Garrel, C.; Besson, G., The role of oxidative stress in amyotrophic lateral sclerosis and Parkinson’s disease. Neurochem. Res. 2010, 35 (10), 1530–1537.

12. Pacher, P.; Beckman, J. S.; Liaudet, L., Nitric oxide and peroxynitrite in health and disease. Physiol. Rev. 2007, 87 (1), 315–424.

13. Mallozzi, C.; Di Stasi, M. A.; Minetti, M., Peroxynitrite-dependent activation of src tyrosine kinases lyn and hck in erythrocytes is under mechanistically different pathways of redox control. Free Radic. Biol. Med. 2001, 30 (10), 1108–1117.

14. Klotz, L. O.; Schroeder, P.; Sies, H., Peroxynitrite signaling: receptor tyrosine kinases and activation of stress-responsive pathways. Free Radic. Biol. Med. 2002, 33 (6), 737–743.

15. Prolo, C.; Álvarez, M. N.; Radi, R., Peroxynitrite, a potent macrophagelderived oxidizing cytotoxin to combat invading pathogens. BioFactors 2014, 40 (2), 215–225.

16. Marla, S. S.; Lee, J.; Groves, J. T., Peroxynitrite rapidly permeates phospholipid membranes. Proc. Natl. Acad. Sci. U.S.A. 1997, 94 (26), 14243–14248.

17. Chen, X.; Chen, H.; Deng, R.; Shen, J., Pros and cons of current approaches for detecting peroxynitrite and their applications. Biomed J 2014, 37 (3), 120–126.

18. Peng, T.; Wong, N. K.; Chen, X.; Chan, Y. K.; Ho, D. H.; Sun, Z.; Hu, J. J.; Shen, J.; El-Nezami, H.; Yang, D., Molecular imaging of peroxynitrite with HKGreen-4 in live cells and tissues. J. Am. Chem. Soc. 2014, 136 (33), 11728–11734.

19. Sun, X.; Xu, Q.; Kim, G.; Flower, S. E.; Lowe, J. P.; Yoon, J.; Fossey, J. S.; Qian, X.; Bull, S. D.; James, T. D., A water-soluble boronate-based fluorescent probe for the selective detection of peroxynitrite and imaging in living cells. Chem. Sci. 2014, 5 (9), 3368–3373.

20. Chen, Z.; Truong, T. M.; Ai, H. W., Development of Fluorescent Probes for the Detection of Peroxynitrite. In Peroxynitrite Detection in Biological Media: Challenges and Advances, 2015; p 186.

21. Ambikapathi, G.; Kempahanumakkagari, S. K.; Lamani, B. R.; Shivanna, D. K.; Maregowda, H. B.; Gupta, A.; Malingappa, P., Bioimaging of Peroxynitrite in MCF-7 Cells by a New Fluorescent Probe Rhodamine B Phenyl Hydrazide. J. Fluoresc. 2013, 23 (4), 705–712.

22. Zielonka, J.; Sikora, A.; Hardy, M.; Joseph, J.; Dranka, B. P.; Kalyanaraman, B., Boronate probes as diagnostic tools for real time monitoring of peroxynitrite and hydroperoxides. Chem. Res. Toxicol. 2012, 25 (9), 1793–1799.

23. Lippert, A. R.; De Bittner, G. C. V.; Chang, C. J., Boronate Oxidation as a Bioorthogonal Reaction Approach for Studying the Chemistry of Hydrogen Peroxide in Living Systems. Acc. Chem. Res. 2011, 44 (9), 793–804.

24. Chen, Z. J.; Ren, W.; Wright, Q. E.; Ai, H. W., Genetically encoded fluorescent probe for the selective detection of peroxynitrite. J. Am. Chem. Soc. 2013, 135 (40), 14940–14943.

25. Chen, Z. J.; Tian, Z.; Kallio, K.; Oleson, A. L.; Ji, A.; Borchardt, D.; Jiang, D. E.; Remington, S. J.; Ai, H. W., The N-B Interaction through a Water Bridge: Understanding the Chemoselectivity of a Fluorescent Protein Based Probe for Peroxynitrite. J. Am. Chem. Soc. 2016, 138 (14), 4900–4907.

26. Chen, Z.; Zhang, S.; Li, X.; Ai, H. W., A high-performance genetically encoded fluorescent biosensor for imaging physiological peroxynitrite. Cell Chem. Biol. 2021, 28 (11), 1542–1553.

27. Liu, C. C.; Schultz, P. G., Adding new chemistries to the genetic code. Annu Rev Biochem 2010, 79, 413-444.

28. Chen, S.; Chen, Z. J.; Ren, W.; Ai, H. W., Reaction-based genetically encoded fluorescent hydrogen sulfide sensors. J. Am. Chem. Soc. 2012, 134 (23), 9589–9592.

29. Zhao, Y.; Araki, S.; Wu, J.; Teramoto, T.; Chang, Y. F.; Nakano, M.; Abdelfattah, A. S.; Fujiwara, M.; Ishihara, T.; Nagai, T.; Campbell, R. E., An expanded palette of genetically encoded Ca2+ indicators. Science 2011, 333 (6051), 1888–1891.

30. Fan, Y.; Chen, Z.; Ai, H. W., Monitoring redox dynamics in living cells with a redox-sensitive red fluorescent protein. Anal. Chem. 2015, 87, 2802–2810.

31. Young, T. S.; Ahmad, I.; Yin, J. A.; Schultz, P. G., An enhanced system for unnatural amino acid mutagenesis in E. coli. J. Mol. Biol. 2010, 395 (2), 361–374.

32. Brustad, E.; Bushey, M. L.; Lee, J. W.; Groff, D.; Liu, W.; Schultz, P. G., A genetically encoded boronate-containing amino acid. Angew. Chem. Int. Ed. Engl. 2008, 47 (43), 8220-8223.

33. Miyawaki, A.; Shcherbakova, D. M.; Verkhusha, V. V., Red fluorescent proteins: chromophore formation and cellular applications. Curr. Opin. Struct. Biol. 2012, 22 (5), 679–688.

34. Martin-Romero, F. J.; Gutierrez-Martin, Y.; Henao, F.; Gutierrez-Merino, C., Fluorescence measurements of steady state peroxynitrite production upon SIN-1 decomposition: NADH versus dihydrodichlorofluorescein and dihydrorhodamine 123. J Fluoresc 2004, 14 (1), 17–23.

35. Emsley, P.; Cowtan, K., Coot: model-building tools for molecular graphics. Acta Crystallogr. Sect. D. Biol. Crystallogr. 2004, 60 (12), 2126–2132.

36. Murshudov, G. N.; Skubák, P.; Lebedev, A. A.; Pannu, N. S.; Steiner, R. A.; Nicholls, R. A.; Winn, M. D.; Long, F.; Vagin, A. A., REFMAC5 for the refinement of macromolecular crystal structures. Acta Crystallogr. Sect. D. Biol. Crystallogr. 2011, 67 (4), 355–367.

37. Krieger, E.; Koraimann, G.; Vriend, G., Increasing the precision of comparative models with YASARA NOVA—a selflparameterizing force field. Proteins: Struct. Funct. Genet. 2002, 47 (3), 393–402.

38. Maier, J. A.; Martinez, C.; Kasavajhala, K.; Wickstrom, L.; Hauser, K. E.; Simmerling, C., ff14SB: Improving the Accuracy of Protein Side Chain and Backbone Parameters from ff99SB. J. Chem. Theory Comput. 2015, 11 (8), 3696–3713.

39. Niu, W.; Guo, J., Expanding the chemistry of fluorescent protein biosensors through genetic incorporation of unnatural amino acids. Mol. Biosyst. 2013, 9 (12), 2961–2970.

40. Liu, X.; Li, J.; Hu, C.; Zhou, Q.; Zhang, W.; Hu, M.; Zhou, J.; Wang, J., Significant expansion of the fluorescent protein chromophore through the genetic incorporation of a metal-chelating unnatural amino acid. Angew. Chem. Int. Ed. Engl. 2013, 52 (18), 4805–4809.

41. Zhang, S.; Ai, H. W., A general strategy to red-shift green fluorescent protein-based biosensors. Nat. Chem. Biol. 2020, DOI: 10.1038/s41589-41020-40641-41587.

42. Akerboom, J.; Carreras Calderón, N.; Tian, L.; Wabnig, S.; Prigge, M.; Tolö, J.; Gordus, A.; Orger, M. B.; Severi, K. E.; Macklin, J. J., Genetically encoded calcium indicators for multi-color neural activity imaging and combination with optogenetics. Front. Mol. Neurosci. 2013, 6, 2.

43. Zhang, J.; Li, Z.; Pang, Y.; Fan, Y.; Ai, H. W., Genetically Encoded Boronolectin as a Specific Red Fluorescent UDP-GlcNAc Sensor. bioRxiv, 2023, doi: https://doi.org/10.1101/2023.03.01.530644.

44. Kabsch, W., Xds. Acta Crystallogr. Sect. D. Biol. Crystallogr. 2010, 66 (2), 125–132.

45. Evans, P. R., An introduction to data reduction: space-group determination, scaling and intensity statistics. Acta Crystallogr. D Biol. Crystallogr. 2011, 67 (Pt 4), 282–292.

46. Evans, P. R.; Murshudov, G. N., How good are my data and what is the resolution? Acta Crystallogr. D Biol. Crystallogr. 2013, 69 (Pt 7), 1204–1214.

47. French, S.; Wilson, K., On the treatment of negative intensity observations. Acta Crystallographica Section A: Crystal Physics, Diffraction, Theoretical and General Crystallography 1978, 34 (4), 517–525.

48. McCoy, A. J.; Grosse-Kunstleve, R. W.; Adams, P. D.; Winn, M. D.; Storoni, L. C.; Read, R. J., Phaser crystallographic software. J. Appl. Crystallogr. 2007, 40 (4), 658–674.

49. Laskowski, Roman A., and Mark B. Swindells. LigPlot+: Multiple ligand–protein interaction diagrams for drug discovery. J. Chem. Inf. Model. 2011, 51 (10), 2778–2786.

50. Dewar, M. J. S.; Zoebisch, E. G.; Healy, E. F.; Stewart, J. J. P., Development and use of quantum mechanical molecular models. 76. AM1: a new general purpose quantum mechanical molecular model. J. Am. Chem. Soc. 1985, 107 (13), 3902–3909.

