## Supporting Information for "Development, Characterization, and Structural Analysis of a Genetically Encoded Red Fluorescent Peroxynitrite Biosensor"

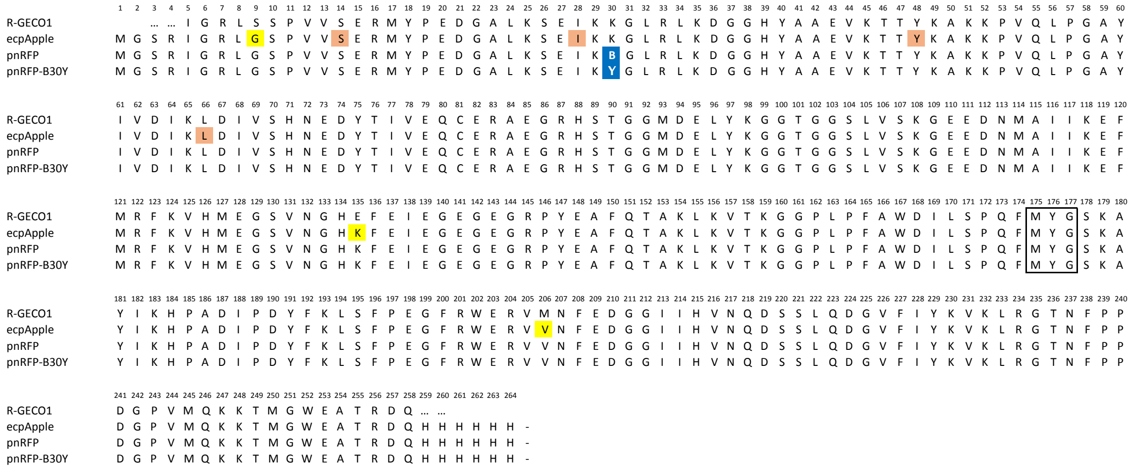


**Fig. S1. Sequence alignment of pnRFP with several relevant variants.** Shaded in yellow are mutations in ecpApple obtained through directed evolution from the circularly permuted RFP fragment in R-GECO1. Shaded in blue is residue 30 in pnRFP or pnRFP-B30Y, and B denotes *p*BoF. Shaded in orange are other residues in ecpApple examined for the genetic incorporation of *p*BoF in this study. Highlighted in the black box are residues 175-177, which are responsible for the formation of the chromophore in each protein.

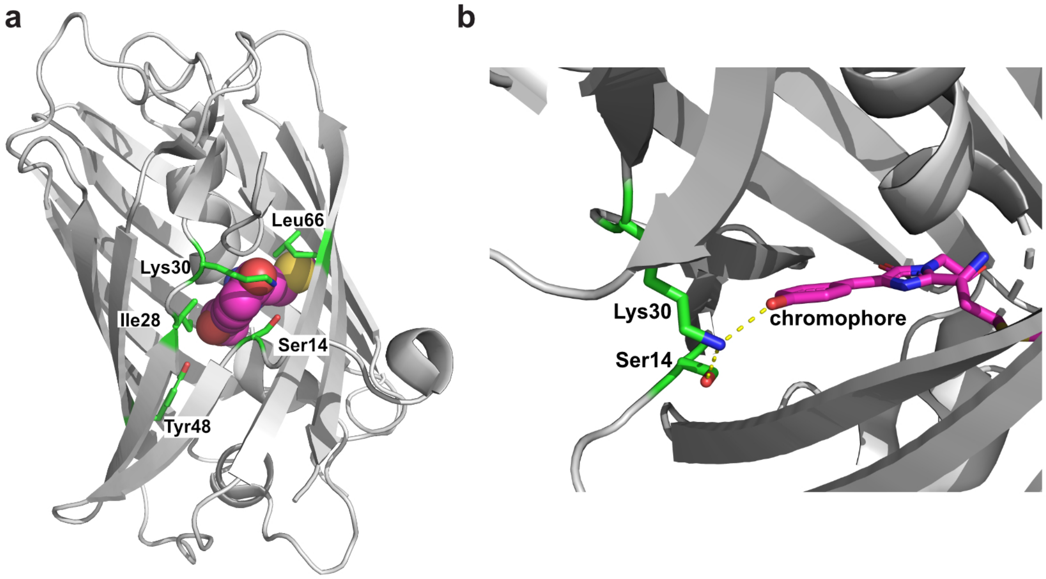


**Fig. S2. Illustration of residues in cpmApple targeted for site-specific incorporation of *p*BoF.** (**a**) Presented in green sticks are residues 14, 28, 30, 48, and 66. The chromophore is shown in magenta balls. (**b**) A detailed illustration of residues 14 and 30 (green sticks) in relation to the chromophore (magenta sticks). H-bonds between them are presented as yellow dashes. These graphs were generated based on the X-ray crystal structure of R-GECO1 (Protein Data Bank (PDB) entry 4I2Y).

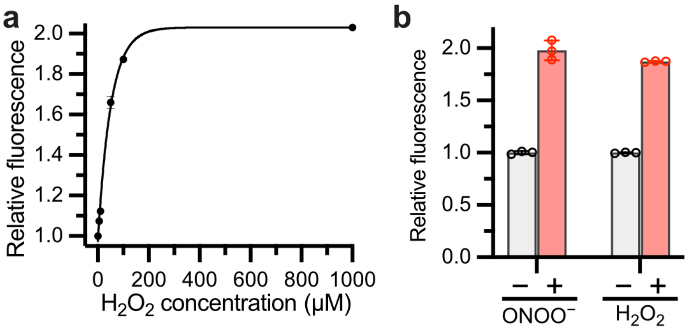


**Fig. S3. Fluorescence responses of ecpApple-S14B.** (**a**) Fluorescence increase of ecpApple-S14B after incubation with the indicated concentrations of hydrogen peroxide for 20 min, showing the sensitivity of ecpApple-S14B to micromolar hydrogen peroxide. (**b**) Comparison of the responses of ecpApple-S14B to 100 µM peroxynitrite or 100 µM hydrogen peroxide. Data are presented as mean ± s.d. of three technical replicates.

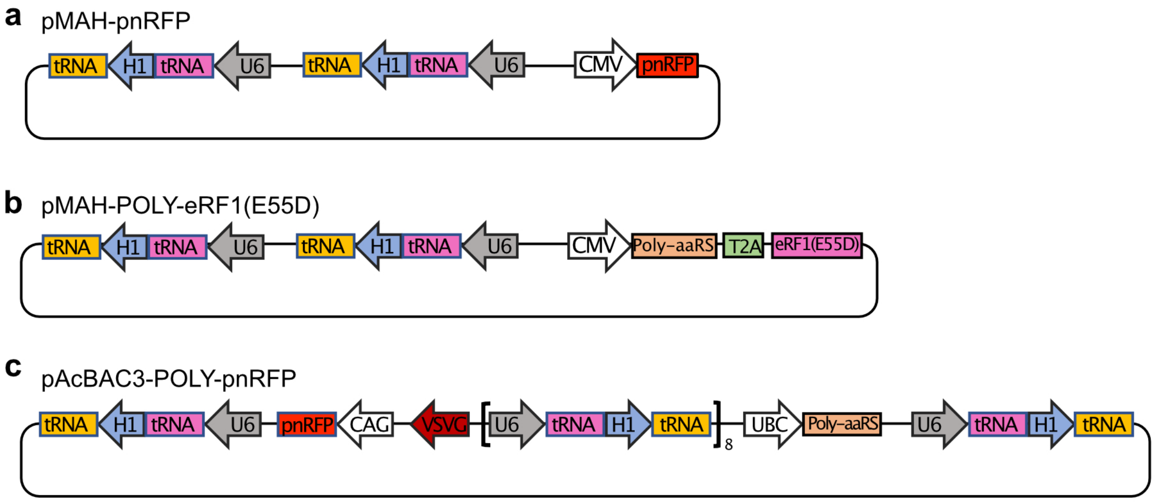


**Fig. S4. Illustration of main genetic elements of the indicated plasmids.** Plasmid maps for pMAH-pnRFP (**a**), pMAH-POLY-eRF1(E55D) (**b**), and pAcBAC3-POLY-pnRFP (**c**). H1 and U6 are promoters to drive the expression of the amber suppression tRNAs. CMV, CAG and UBC are promoters to drive the expression of the orthogonal aminoacyl-tRNA synthetase (Poly-aaRS), the pnRFP sensor, or a release factor mutant (eRF1(E55D)) that competes with endogenous eRF1 to reduce amber codon termination. pAcBAC3 was initially designed for packing baculovirus, but this study uses it as a plasmid vector for transient transfection.

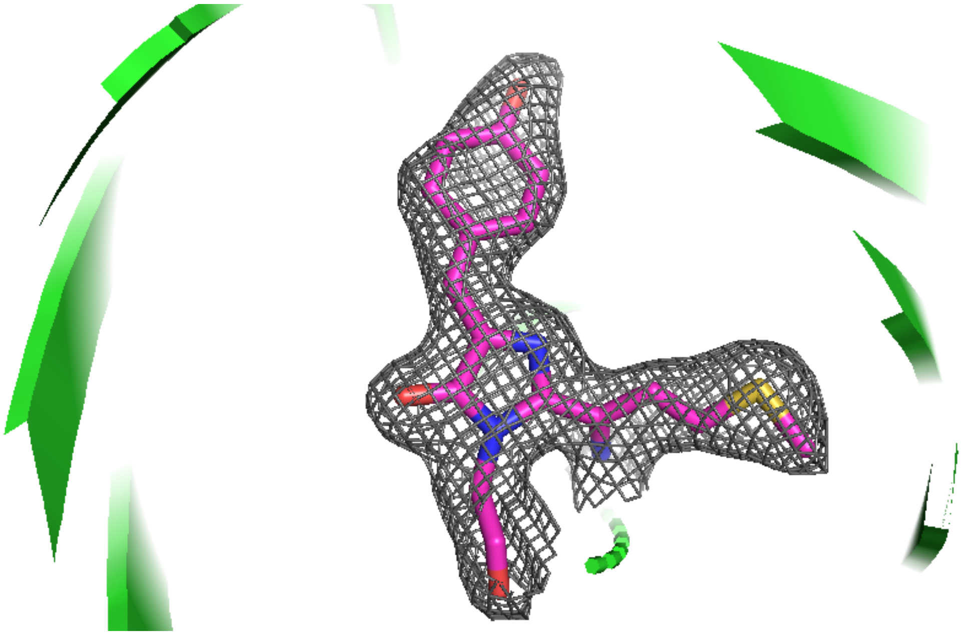

**Fig. S5. Fitting of the chromophore of pnRFP in the 2Fo-Fc electron density map at 1.0σ.** The chromophore adopts a clear *cis* conformation.

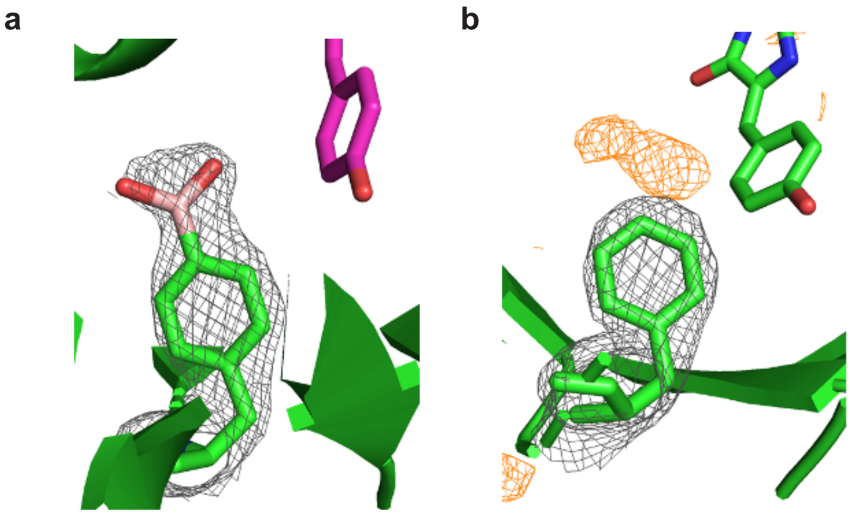


**Fig. S6. Conformation analysis of *p*BoF30 in pnRFP.** (**a**) The 2Fo-Fc electron density map (grey mesh) well describes *p*BoF30 at 1.0 σ. (**b**) After substituting *p*BoF with Phe, a positive Fo-Fc difference electron density map (orange) was shown on the top of Phe at 3.0 σ, demonstrating a missing portion of the residue.

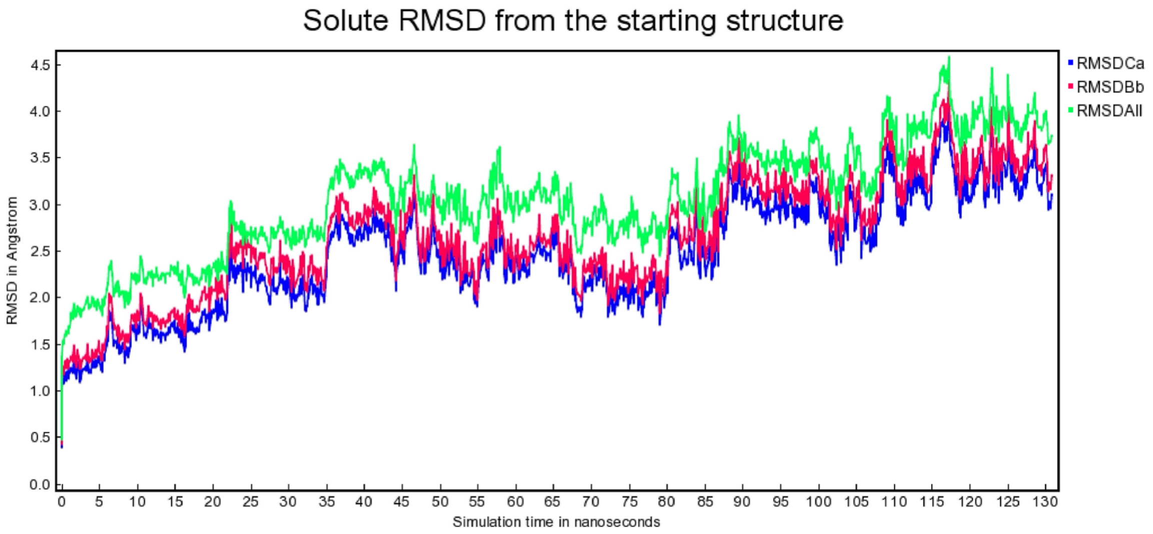


**Fig. S7. MD simulation trajectory of ecpApple-S14B under the AMBER14 force field in YASARA.** The trajectory indicates that the simulated model was in the equilibrium state.

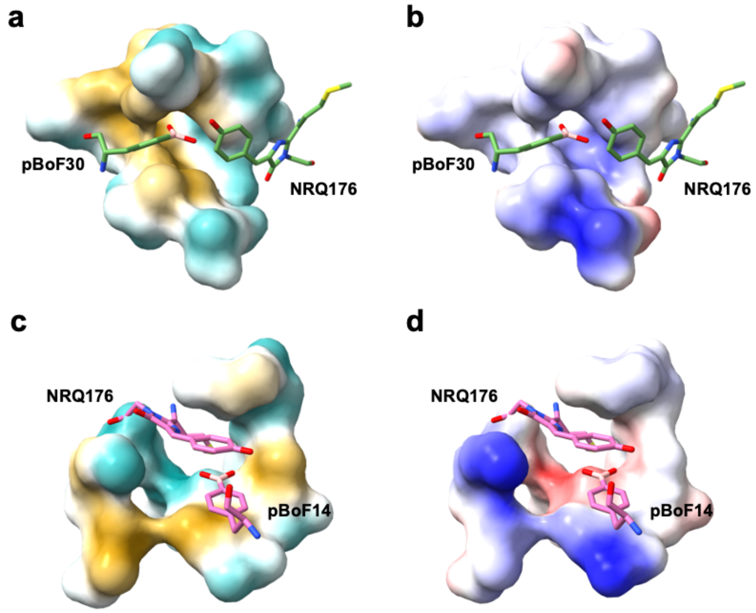


**Fig. S8. Hydrophobicity and electrostatic potentials of residues surrounding *p*BoF in pnRFP (a, b) and ecpApple-S14B (c, d).** In panels a and c, hydrophobicity surface was mapped according to the hydrophobicity scale of Kyte and Doolittle (1) and colored from cyan for the most hydrophilic, to white, to deep sand for the most hydrophobic. In panels b and d, electrostatic potentials of the surrounding residues were calculated by Coulomb's law (2), and the surface was colored with the calculated values from blue for the most positive potential, to white, to red for the most negative potential. Compared with *p*BoF30 in pnRFP, the boronic acid-head group of *p*BoF14 in ecpApple-S14B is in a more hydrophilic and more negatively charged environment. The surface coloring and visualization of the models were achieved with ChimeraX (3).

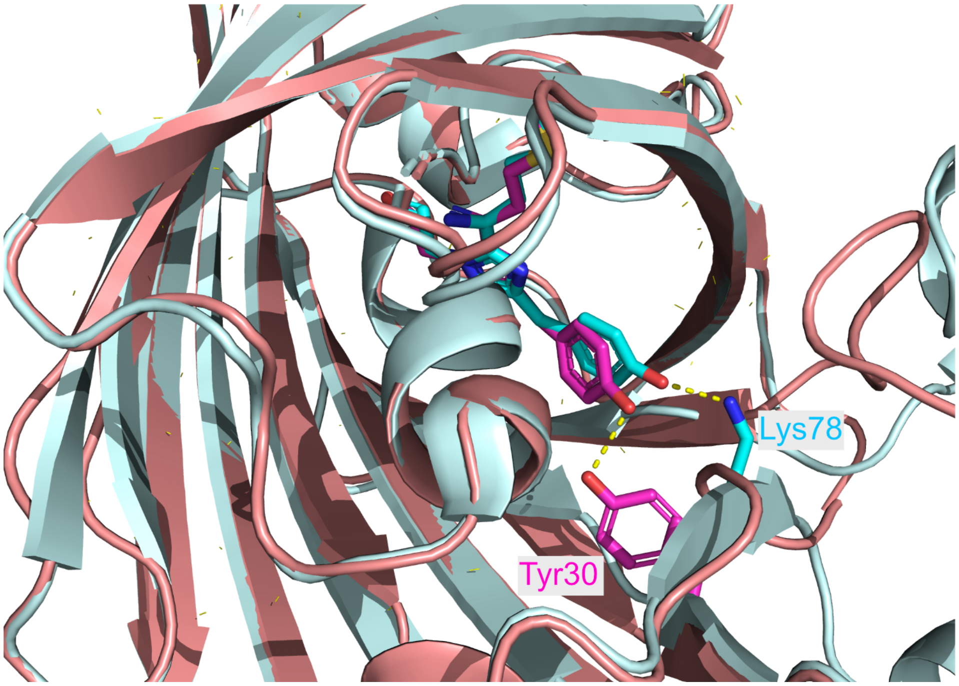


**Fig. S9. Overlay of the crystal structures of R-GECO1 (cyan) and pnRFP-B30Y (pink/magenta).** The locations of Lys78 of R-GECO1 and Tyr30 of pnRFP-B30Y in relation to the chromophores are highlighted.

**Table S1. X-ray crystallography data collection and structure refinement statistics.**

| | pnRFP<br>PDB 7LQO | pnRFP-B30Y<br>PDB 7LUG |
|---|---|---|
| *Data collection* | | |
| Space group | C121 | C121 |
| Unit cell dimensions | | |
| a, b, c (Å) | 84.20, 34.57, 88.81 | 84.28, 34.11, 89.22 |
| A, β, γ (°) | 90.00, 110.90, 90.00 | 90.00, 110.79, 90.00 |
| Resolution range (Å) | 41.81 - 2.10 (2.16 - 2.10) | 41.85 - 1.95 (2.00 - 1.95) |
| Total reflections | 50624 (2135) | 62455 (4587) |
| Unique reflections | 13553 (831) | 17539 (1245) |
| Completeness (%) | 94.9 (70.3) | 99.2 (99.5) |
| I/δ (I) | 13.3 (1.78) | 10.8 (2.57) |
| $R_{merge}$ (%) | 0.052 (0.433) | 0.058 (0.445) |
| CC(1/2) | 0.998 (0.805) | 0.998 (0.814) |
| *Refinement* | | |
| Resolution range (Å) | 41.81 – 2.10 | 41.85 – 1.95 |
| No. of Reflections | 12873 | 16652 |
| No. of non-hydrogen atoms | 1933 | 1903 |
| No. of waters | 51 | 65 |
| $R_{work}$/ $R_{free}$ | 0.188/0.254 | 0.189/0.212 |
| RMS bond length (Å) | 0.0080 | 0.0050 |
| RMS bond angles (°) | 1.571 | 1.531 |
| Average B value (Å$^2$) | 38.0 | 34.0 |
| *Ramachandran plot* | | |
| Favored (%) | 97.79 | 99.11 |
| Allowed (%) | 2.21 | 0.89 |
| Outliers (%) | 0.00 | 0.00 |

**Table S2. The sequences of oligos used in this work.**

| Oligo Name | Sequence (5′→3′) |
|---|---|
| cpRFP_F | GGAATTAACCATGGGCTCGAGAATAGGTCGGCTGGGCTCA |
| cpRFP_R | TCCGCCAAAACAGCCAAGCTTAATGATGGTGGTGATGGTG |
| ecpApple-S14TAG-F | ATAGGTCGGCTGGGCTCACCCGTAGTTTAGGAGCGGATGTACCCCGAGG |
| ecpApple-I28TAG-F | GCGAGTAGAAGAAGGGGCTGAGGCTGAA |
| ecpApple-I28TAG-R | CAGCCCCTTCTTCTACTCGCTCTTCAG |
| ecpApple-K30TAG-F | GCGAGATCAAGTAGGGGCTGAGGCTGAA |
| ecpApple-K30TAG-R | CAGCCCCTACTTGATCTCGCTCTTCA |
| ecpApple-Y48TAG-F | CCTAGAAGGCCAAGAAGCCCGTGCAGCTGCCCGGC |
| ecpApple-Y48TAG-R | GGGCTTCTTGGCCTTCTAGGTGGTCTTGAC |
| ecpApple-L66TAG-F | ATCAAGTAGGACATCGTGTCCCACAAC |
| ecpApple-L66TAG-R | GTGGGACACGATGTCCTACTTGATGTCGACGATGTA |
| ecpApple-Y176TAG-F | TCCCCTCAGTTCATGTAGGGCTCCAAGGCCTACATT |
| ecpApple-Y176TAG-R | GGAGCCCTACATGAACTGAGGGGACAGGATGTC |
| pnRFP_F | CCAGGTCCAACTGCACGGAAGCTTGCCACCATGGATAGGTCGGCTGGGC TC |
| pnRFP_R | TCAGCGGGTTTAAACGGGCCCTTAATGATGGTGGTGATGGTGTTGGTCA CGCGTAGCC |
| pBac3-F | GGAGGCCACCATGGGCTCGAGAATAGGTCGGCTGGGCTCA |
| pBac3-R | TCGACTTAACGCGTTGAATTCATGCATTTAATGATGGTGGT |

### References

1. J. Kyte, R. F. Doolittle, A simple method for displaying the hydropathic character of a protein. *J. Mol. Biol.* **157**, 105-132 (1982).
2. The detail on Coulomb's law applied in the calculation is presented on the following website: https://www.cgl.ucsf.edu/chimera/data/downloads/1.8/docs/ContributedSoftware/coulombic/coulombic.html.
3. E. F. Pettersen *et al.*, UCSF ChimeraX: Structure visualization for researchers, educators, and developers. *Protein Sci.* **30**, 70-82 (2021).
